# Arctic-Atlantic gradient shaped PFAS exposure variability in sympatric guillemot species off Iceland

**DOI:** 10.1101/2025.09.15.676265

**Authors:** Rui Shen, Ralf Ebinghaus, Daniel Giddings Vassão, Norman Ratcliffe, Thomas Larsen

## Abstract

Per- and polyfluoroalkyl substances (PFAS) are persistent organic pollutants of growing environmental concern in marine ecosystems. While previous approaches have focused on mean concentrations, we here propose treating PFAS exposure variability as an ecological signal rather than statistical noise. To examine this variability-as-signal hypothesis, we analysed PFAS concentrations in plasma and stable isotopes in plasma and red blood cells from 112 individuals of two sympatric guillemot species (*Uria lomvia*, n = 45; *U. aalge*, n = 67) across five Icelandic colonies during the 2018 breeding season. The dual-tissue isotopic approach allowed us to assess foraging consistency across different temporal scales, providing context for interpreting PFAS exposure patterns. PFAS variability was dominated by two compound groups: Long-chain perfluoroalkyl carboxylic acids (PFCAs, 79% of variance) and perfluorooctane sulfonate (PFOS, 13% of variance). We standardised individual variability into Z-scores to quantify individual expression of these exposure patterns, revealing three distinct variability clusters corresponded with oceanographic transitions between Arctic and Atlantic waters. Segmented regression analysis showed that significant threshold effects at the Arctic-Atlantic habitats (distinguished by isotopic breakpoints δ^13^C_consist_ = 0.19, δ^15^N_consist_ = 0.00) with contrasting ecological drivers: PFOS variability responded to habitat-driven shifts (δ^13^C, *p* < 0.01) and PFCA variability to trophic indicators (δ^15^N). Both species exhibited similar PFAS patterns when foraging in the same water masses, with notable exceptions where niche partitioning occurred at oceanographic boundaries. Our findings demonstrate that water mass characteristics and foraging strategies create structured PFAS variability patterns that reflect local ecological adaptations within broader geographical gradients. This variability-focused framework reveals ecological dimensions of contamination that complement traditional mean-based approaches and may improve understanding of contaminant risks in rapidly changing marine ecosystems.

## Introduction

Marine ecosystems, encompassing more than 90% of Earth’s habitable biosphere volume^1^, are threatened by multiple anthropogenic pressures^2^, including the ubiquitous presence of synthetic chemical compounds such as per- and polyfluoroalkyl substances (PFAS)^3^. These persistent organic pollutants, notable for their stable carbon-fluorine bonds, are exceptionally resistant to environmental degradation processes^4^ and accumulate in living organisms^5, 6^. The toxic effects of PFAS pose measurable risks to both human and ecosystem health, especially within marine environments where biomagnification can be pronounced through efficient trophic transfer from phytoplankton to apex predator^6–9^.

The unique physicochemical properties of PFAS determine their bioaccumulation patterns in marine organisms.^10, 11^ Marine predator tissues show a predominance of perfluorooctane sulfonate (PFOS) and long-chain perfluoroalkyl carboxylic acids (PFCAs).^7, 12–14^ PFOS demonstrates enhanced protein-binding affinity and prolonged retention in organisms due to its sulfonate group, which forms stronger bonds with serum proteins than the carboxylate groups of PFCAs.^15–18^ These molecular-level differences create distinct exposure patterns in predators, with PFOS exhibiting greater persistence and integrating exposure over longer time periods compared to PFCAs of similar chain length.^11, 19^ Such compound-specific dynamics fundamentally shape how PFAS are transferred and magnified across trophic levels in marine food webs.^6, 8, 20, 21^

PFAS bioaccumulation research has relied heavily on mean-based metrics like trophic magnification factors (TMFs) and biomagnification factors (BMFs).^20–25^ These approaches obscure ecological nuances by averaging exposure across temporal and spatial scales.^21^ Rather than dismissing variability as statistical noise, we propose recognising it as biologically informative. Variability in PFAS in tissue concentrations within and among populations represent ecological signals that reveal contaminant pathways that mean-based analyses may not capture. This reframing turns individual differences from mere measurement error into ecological insight. For instance, when seabirds display constrained variability, it may indicate specialised foraging within consistent habitats and prey, whereas high variability among individuals can reflect exposure to seasonal prey communities or spatially heterogenous habitats.^26, 27^ Recognising these variability pattern becomes ever more critical as climate change reshapes marine ecosystems, forcing predators to shift their foraging behaviour and, in turn, modifying contaminant exposure pathways.

Spatial patterns of PFAS distribution in marine ecosystems reflect the dynamic integration between ecological and oceanographic processes.^28^ Previous research has primarily attributed these patterns to long--range transport and geographic variability.^29–33^

However, distribution gradients also stem from differences in food web structures and habitat dynamics.^34^ In Arctic waters, PFAS exposure tends to be relatively homogeneous within species because individuals are constrained to narrow, well-segregated habitats and rely on similar, energy-rich prey, leading to reduced inter-individual differences in contaminant uptake. These marine systems typically display more modular and less connected food web structures than temperate regions.^26, 35^ Atlantic waters, conversely, support more diverse prey assemblages including various forage fish and mesopelagic fauna. This greater connectivity and year-round prey availability may create more integrated trophic networks with higher species interactions and diverse feeding opportunities compared to Arctic systems.^36^

Seabirds, as central-place foragers during breeding seasons, provide an ideal model to explore these dynamics.^37, 38^ The period offers a natural framework for examining PFAS exposure variability, as central-place foraging behaviour confines individuals to localised ranges for chick provisioning^39–42^. As a result, central-place foraging itself can promote more homogeneous exposure patterns among individuals, an effect that is independent of whether birds breed in Arctic or temperate environments.^43, 44^ Case in point, two sympatric seabird species, Brünnich’s guillemots (*Uria lomvia*; UL) and common guillemots (*Uria aalge*; UA), have overlapping ecological niches despite UL’s preference for more lipid-rich prey and colder foraging habitats.^45^ In contrast, outside the breeding season, both species disperse across diverse habitats and prey communities^46 47^, potentially leading to greater heterogeneity in contaminant exposure. By focusing on the breeding season of these two species, this study leverages the ecological consistency of central-place foraging to disentangle how spatial habitat use and trophic interactions shape PFAS exposure patterns.

Iceland’s coastal waters, situated at the confluence of cold Arctic and warm Atlantic currents, present a unique natural laboratory for investigating PFAS exposure in marine ecosystems.^48–50^ This region lies at the boundary between Arctic and temperate waters, creating distinct oceanographic and ecological zones.^51^ To the north, cold Arctic waters with seasonal sea ice coverage generally exhibit lower primary productivity than temperate waters, except for localized hotspots such as ice edges and polynyas.^51, 52^ These Arctic systems experience pronounced seasonal production limited to a one to four month period during spring and summer^53^, creating more dynamic trophic relationships^54^. In contrast, the warmer Atlantic-influenced waters to the south without sea ice, generally support higher and more consistent primary productivity.^51, 55^ Atlantic influenced ecosystems around Iceland have more complex, stable, and highly connected food webs than Arctic systems, because the mixing of Arctic and Atlantic water masses introduces a wider range of prey species and ecological niches.^36, 56^ This convergence creates abrupt ecological transitions, where isotopic baselines (δ^13^C, δ^15^N) and prey communities shift sharply across water masses, which in turn makes it possible to test how oceanographic thresholds structure PFAS variability patterns.

To gain insight into foraging consistency at individual levels, we employ stable isotope analysis across two blood fractions with different turnover rates. Plasma integrate dietary information over short periods (~1 week), and red blood cells reflect longer timeframes (~3 – 4 weeks).^57^ By integrating isotopic values across these tissues, we assess how temporal consistency in foraging behaviour shape individual PFAS exposure patterns. This dual-tissue approach provides a more detailed understanding of how individual foraging strategies translate into PFAS variability patterns than is possible with single-tissue analysis.

In this study, we explore contaminant dynamics in marine ecosystems by reconceptualising PFAS exposure variability as an ecological signal rather than statistical noise. Using Iceland’s position at the Arctic-Atlantic interface, we hypothesize that PFAS exposure variability contains structured ecological signals that reflect species-specific foraging strategies and oceanographic influences. Specifically, we predict that UL, with their preference for Arctic waters and more specialized foraging will exhibit more homogeneous (constrained) PFAS variability patterns, while UA, with their broader habitat use and more generalist foraging behavior, will show greater inter-individual variability in PFAS exposure patterns.

## Methods and Materials

### Study Design

We analysed 112 blood samples from UA (n = 67) and UL (n = 45) collected at five colonies across Iceland during June and July 2018 (Fig. 1).^58^ The sampling locations encompassed diverse oceanographic influences along both north-south and east-west gradients. Three northernmost colonies (NW, N, NE) harboured both UA and UL populations, while two southernmost colonies (SE, SW) harboured UA only. At the three northernmost colonies where both species coexist, we collected 15 samples per species per colony (n = 45 UL, n = 45 UA from northernmost colonies). At the two southernmost colonies where only UA breeds, we collected 15 samples at SE and 7 samples at SW due to logistical constraints. We separated blood samples into cells and plasma to evaluate guillemot foraging across different temporal scales. We measured PFAS concentrations in plasma, as these compounds exhibit strong binding affinity to albumin, a primary blood plasma protein. Stable isotope patterns in the North Atlantic show distinct spatial trends: δ^13^C values follow a latitudinal gradient, with higher values in Atlantic-influenced waters than in Arctic waters, while δ^15^N values indicate both trophic position and local oceanographic conditions.^59^ The presence of both species at northernmost colonies facilitated direct interspecies comparisons of PFAS exposure. The balanced sampling design at northernmost colonies (n = 15 per species) provides adequate statistical power for direct interspecies comparisons of PFAS exposure, while the presence of both species at these northernmost colonies facilitates examination of species-specific responses to shared oceanographic conditions. Migration patterns differ between species: UL undertake extensive migrations from their high-latitude breeding sites to lower-latitude wintering areas, with the NW colony typically wintering in Southwest Greenland, and N and NE colonies wintering off West Greenland and north of Iceland.^60^ UA generally winter closer to their breeding colonies, though some individuals migrate to the Mid-Atlantic Ridge or the waters between the Faroe Islands, Scotland, and Shetland Islands.^61^

**Figure 1.**
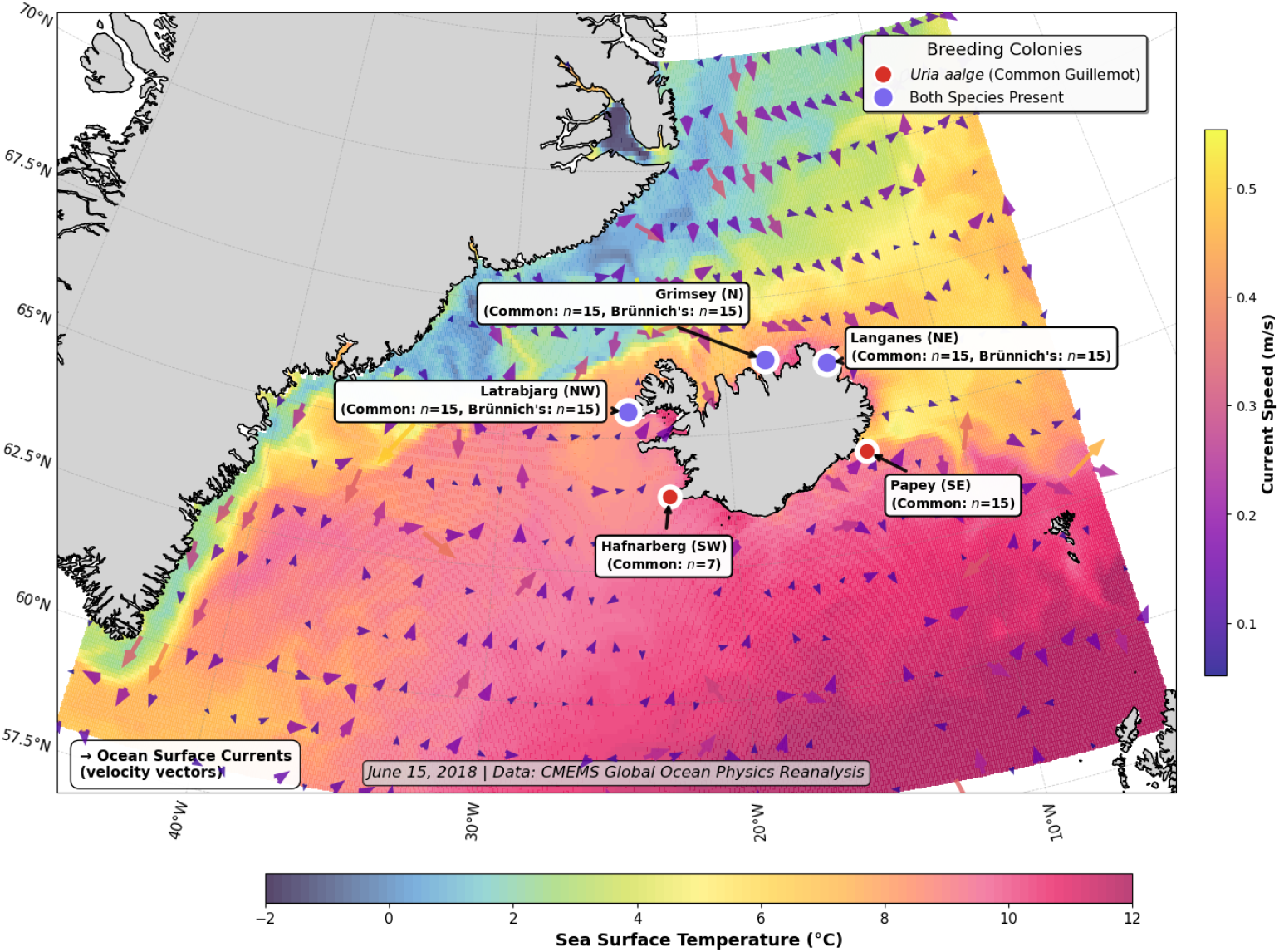
Sea surface temperatures (SSTs) around Iceland in June 2018 highlighting major ocean currents: the warm Irminger and North Atlantic Currents, and the cold East Greenland and East Icelandic Currents. This study sampled guillemots in five colonies. Látrabjarg (NW; 65.50°N, 24.52°W) is influenced by the interface between the warm Irminger Current and the cold East Greenland Current. Grímsey Island (N; 66.57°N, 18.02°W) lies within the warmer waters of the Irminger Current, with some influence from the cooler East Icelandic Current. Langanes peninsula (NE; 66.38°N, 14.54°W) sits where the Irminger and East Icelandic Currents meet. Papey (SE; 64.59°N, 14.18°W) is situated in the cool East Icelandic Current, near its convergence with the warm North Atlantic Current. Hafnaberg (SW; 63.75°N, 22.75°W) is affected by the warmer waters of the North Atlantic Current. Each of the three northern colonies hosted both common (*Uria aalge*, UA) and Brünnich’s guillemots (*Uria lomvia*, UL). NW is the largest colony with 343,900 guillemot pairs (UA:UL ratio 1.9:1) and N the second largest with 71,400 pairs (UA:UL ratio 16.4:1) as per a 2008 census ^62^. Only UA breeds in the two southern colonies, SE and SW.

### Sample Collection

To ensure a representative cross-section of Icelandic seabird populations, all blood samples were collected from selected breeding colonies, as described in Bonnet-Lebrun.^63^ Approximately 1 mL of blood was collected from each seabird via venipuncture, using rapid, minimally invasive sampling techniques to ensure bird welfare. Post-collection, blood samples were centrifuged using a Micro Star 12 centrifuge (VWR, Leuven, Belgium) at 12,300 rpm for 4 minutes to separate plasma and cells. The separated components were transferred to glass petri dishes and dried at ambient temperature in a desiccator for 4 to 6 days (detailed description the Supporting Information).

### PFASs Chemical Analysis

PFAS analysis was performed on plasma samples at Helmholtz-Zentrum Hereon (Geesthacht, Germany). The targeted analysis included 14 PFAS compounds: nine PFCAs (C5 - C13), four

PFSAs (C4, C6, C8, C10), and HFPO-DA (complete list in Table S1, SI).

PFAS extraction from dried plasma samples was performed using a modified quaternary ammonium salt-based ion-pairing method.^64, 65^ Prior to extraction, samples were fortified with ^13^C-labeled PFAS internal standards and buffered with sodium carbonate (1 mL, 0.5 M, pH 10.0). The detailed extraction protocol is provided in the Supporting Information.

Analysis was conducted using HPLC-MS/MS with electrospray ionization in multiple reaction monitoring (MRM) mode. Chromatographic separation was achieved using a Synergi Fusion-RP C18 column with a water-methanol gradient containing ammonium acetate buffer. Method detection limits were 0.05 - 0.07 ng/mL for all target compounds, with data reported on dry weight basis (ng/g DM) consistent with the dried sample preparation approach. Detailed analytical parameters and QA/QC procedures are provided in the SI.

### Stable Isotope Analysis

The isotope data for blood cells reported in this study have already been published by Bonnet-Lebrun.^63^ The plasma isotope data, however, are presented here for the first time. The isotope data are expressed in delta (δ) notation:

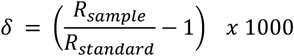

Where *R*_*sample*_ is the ratio of the heavy to light isotope in the sample, and *R*_*standard*_ is the ratio of the heavy to light isotope in the standard. To express the isotopic data as per mil (‰), they are multiplied by 1000. The isotope ratios are expressed relative to Vienna Pee Dee Belemnite (VPDB) for carbon and atmospheric air for nitrogen. Elemental content and bulk isotope values of blood cells were prepared according to Bonnet-Lebrun ^63^ and determined at the Stable Isotope Facility of the Experimental Ecology Group, GEOMAR, Kiel with a customised, high sensitivity elemental analyser connected to a stable isotope ratio mass spectrometer (DeltaPlus Advantage, Thermo Fisher Scientific, Germany) as described by Hansen and Sommer ^66^. The standard deviations for δ^13^C and δ^15^N and ranged from ± 0.15% to ± 0.25% (n = 3).

## Statistical Analysis

All statistical analyses were conducted using Python 3.9.16 with associated libraries, including Pandas 1.5.3, NumPy 1.24.3, SciPy 1.10.1, Scikit-learn 1.2.2, Statsmodels 0.13.5, Matplotlib 3.7.1, and Seaborn 0.12.2. Our analytical approach consisted of four sequential steps designed to characterize PFAS variability patterns and their relationships with ecological indicators.

First, PFAS concentration data were log-transformed to improve normality. We then conducted Principal Component Analysis (PCA) on these log-transformed concentrations to reduce the dimensionality of the multi-compound dataset and identify the primary patterns of covariation among PFAS compounds across individual birds Based on the patterns revealed by PCA (Supplementary Information), we then standardised each individual’s PC scores into Z-scores to quantify how much each bird expresses these covariation patterns. A Z-score of zero represents average variability for that exposure pattern across the population. Negative Z-scores indicate more constrained (consistent) exposure patterns. Positive Z-scores indicate more variable (heterogeneous) exposure patterns. Note that in this study, the application and interpretation of Z-scores is tailored to the specific PCA loading patterns found in this dataset, allowing us to identify individuals with consistently constrained versus heterogeneous exposure patterns that may reflect different foraging strategies. Second, we implemented K-means clustering to characterize population-level patterns in PFAS variability. We determined the optimal number of clusters using silhouette analysis, which maximizes within-cluster similarity and between-cluster separation. Silhouette scores (−1 to +1) measure clustering quality, where higher values indicate better cluster assignment, zeros suggest borderline cases, and negative values indicate likely misclassification. This approach allowed us to identify distinct PFAS variability profiles across the study populations. Finally, to detect potential threshold effects in relationships between isotopic consistency and PFAS variability patterns, we employed bivariate segmented regression. We calculated isotopic consistency scores (δ^13^C_consist_ and δ^15^N_consist_) using PCA to capture the shared consistency between tissues followed by standardisation into Z-scores. A consist score of zero represents the overall population mean, with positive and negative scores indicating consistently higher or lower isotopic values across tissues, respectively. The absolute magnitude of these scores reflects how strongly an individual’s isotopic pattern deviates from the population mean across both temporal scales. For the bivariate segment analysis, the dataset was split into training (80%) and testing (20%) sets. Optimal breakpoints for segmented regression were identified by minimizing the sum of squared residuals. Models included base effects for both isotopic indicators, segment effects for values above identified breakpoints, and interaction terms. Model performance was evaluated using *R*^2^ and RMSE. Based on the identified breakpoints, individuals were classified into habitat-associated segments to assess how oceanographic regimes influenced PFAS variability patterns. All statistical tests were two-tailed, with significance levels set at *α* = 0.05. 95% confidence intervals were reported where appropriate. Detailed descriptions of these calculations and additional statistical methods are provided in the Supplementary Information.

## Results

### PFAS Variability Patterns

Analysis of PFAS concentrations in plasma samples revealed consistent patterns of exposure across the two seabird species UA and UL. Long-chain PFCAs (C9 - C13) and PFOS (linear and branched) were the dominant compounds detected in all samples. PFOS accounted for 62 - 73% of the total PFAS burden in UA and 70 - 89% in UL, highlighting its predominance in both species. Among PFCAs, PFUnDA (C11) and PFTrDA (C13) were the most abundant, with median concentrations ranging from 10.0 to 23.3 ng/g DM in UA and 6.6 to 10.0 ng/g in UL. Short-chain PFCAs (e.g., PFPeA, PFHxA) and PFBS as well as HFPO-DA were not detected in any samples. Detection frequencies and concentration ranges are summarized in Table 1, while Fig. 2 illustrates median PFAS profiles across species and colonies.

**Table 1.**
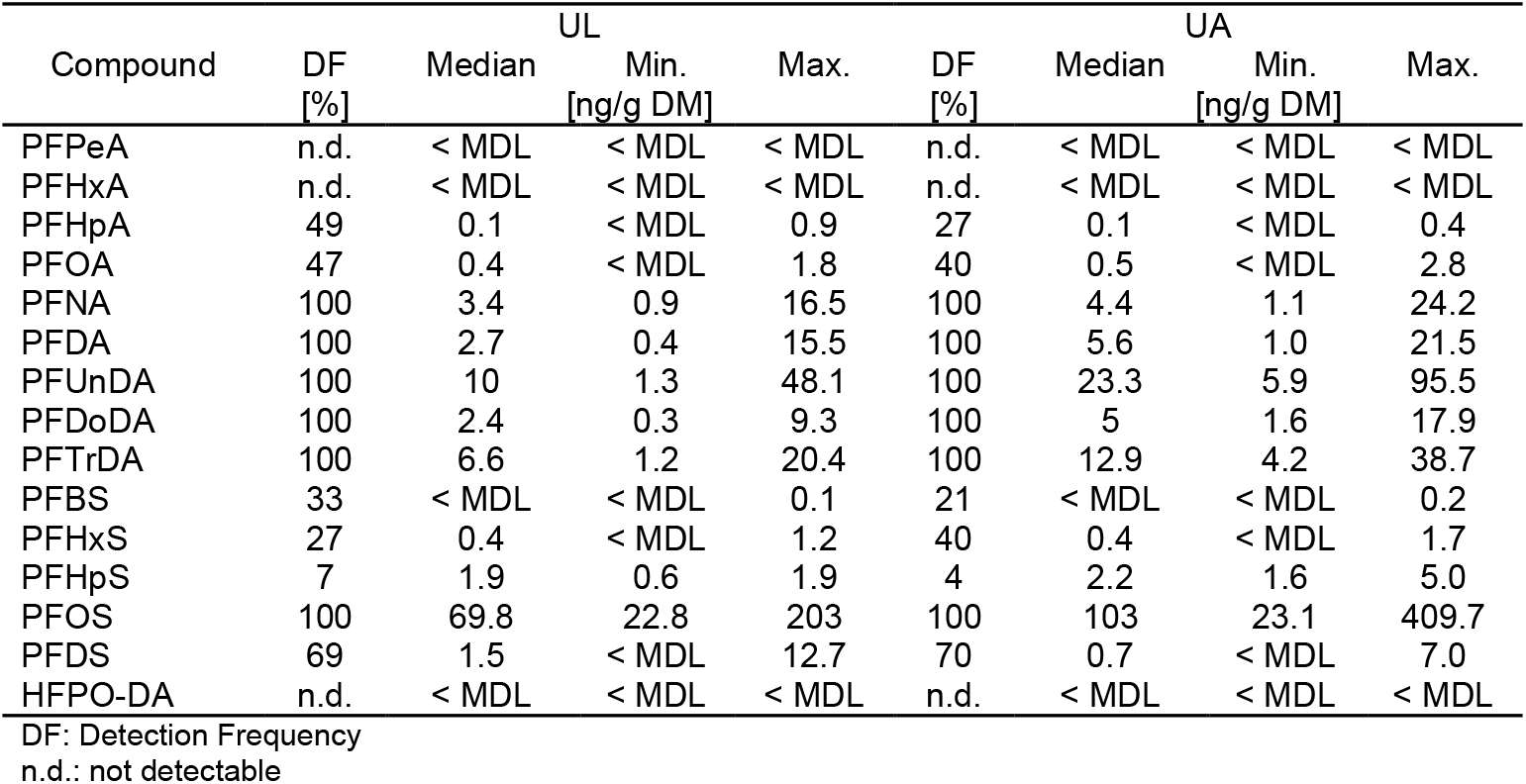
Detection rates and concentration ranges of PFAS compounds in UA and UL samples.

**Figure 2.**
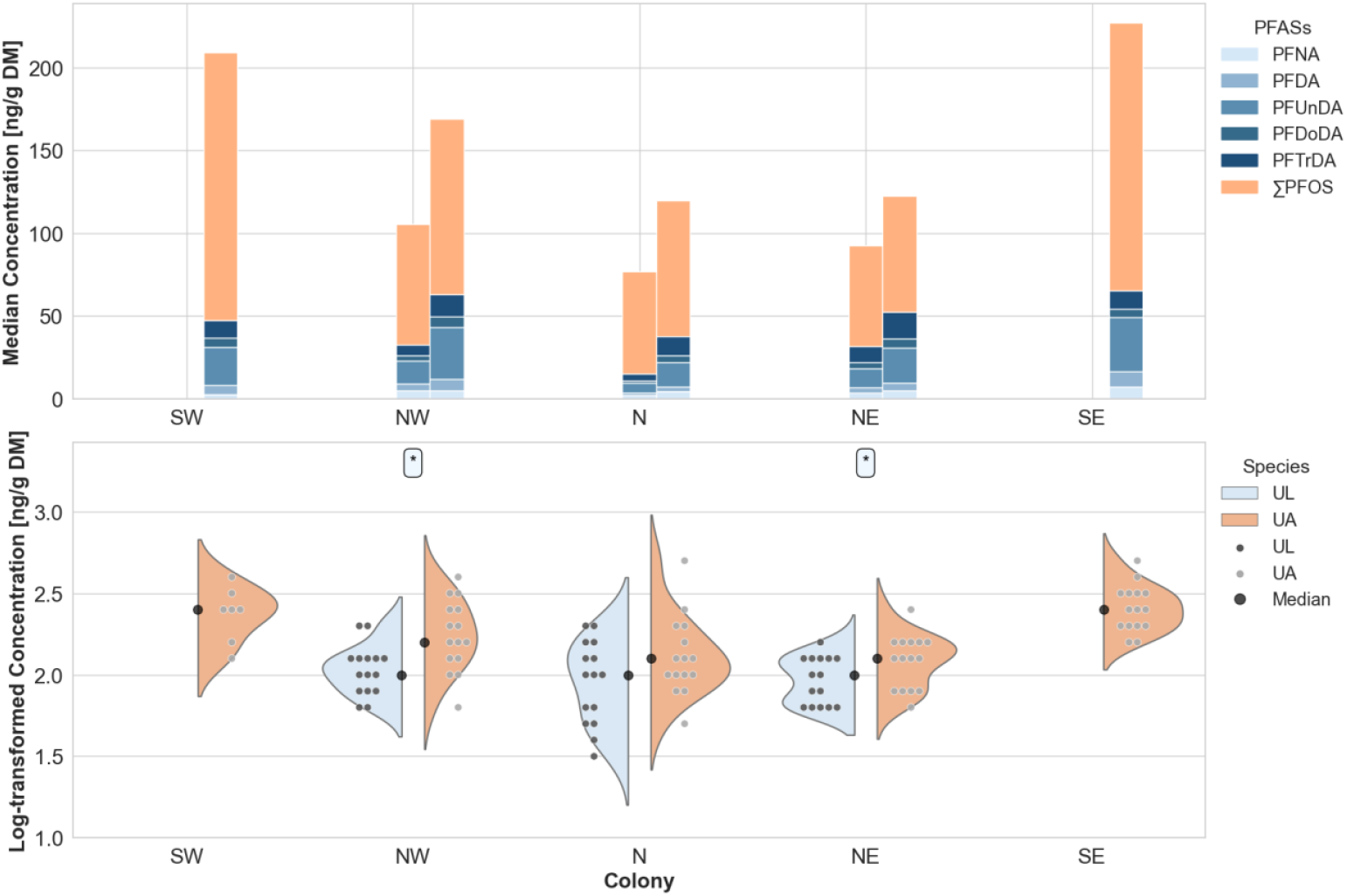
PFAS (per- and polyfluoroalkyl substance) profiles between species (*Uria aalge* (UA) and *Uria lomvia* (UL)) and across their colonies (SW, NW, N, NE, SE) in Iceland. Upper panel: Median PFASs concentrations of five long-chain perfluoroalkyl carboxylic acids (C9-C13 PFCAs) and PFOS (perfluorooctane sulfonate, linear and branched) are shown as stacked bars. Lower panel: Log-transformed PFASs concentrations (including both C9-C13PFCAs and PFOS) are displayed using split-violin plots. Each data point represents an individual bird, while the split-violin shapes reflect the overall distribution for each colony and species. Black dots indicate median values for each species at each colony, and asterisks represent significance levels between species at sympatric colony (* *p* ≤ 0.05).

To identify the underlying exposure patterns driving PFAS variability within individuals, we investigated how PFAS covary using PCA. PCA revealed two main sources of variability in PFAS exposure patterns. The first principal components (PC1), which was mainly driven by long-chain PFCAs (C9 - C13), explained 79% of the variation. The second component (PC2) was driven by PFOS, which varied independently from the other compounds, and accounted for an additional 13% of the variability (detailed results in Supplementary Information, Fig. S1). Based on these distinct patterns, coordinated PFCA behaviour versus independent PFOS behaviour, we applied standardized Z-scores, denoted as Z_PFCA_ (for long-chain PFCAs) and Z_PFOS_ (for PFOS), to measure how much each individual bird expresses these different exposure patterns. Z_PFCA_ ranged from −2.3 to 2.9 across all birds. Northernmost UA showed predominantly negative Z-scores (median: −0.6 to 0.2) with relatively constrained variability. In contrast, northernmost UL exhibited predominantly positive Z-scores (median: 0.3 to 1.1) with greater variability in pattern expression. Z_PFOS_ ranged from −2.1 to 3.4. Southernmost UA showed elevated positive Z-scores (median: 0.7 to 0.8) with high variability in PFOS pattern expression (max values, SW: 3.4; SE:2.7). Northernmost populations of both species showed more constrained PFOS patterns, exhibiting predominantly negative Z-scores across colonies (median, UA: −0.7 to −0.2; UL: −0.5 to 0.4).

To categorize birds according to their PFAS exposure variability profiles, we performed K-means clustering analysis. K-means clustering analysis (k = 3) identified three distinct patterns in PFAS exposure variability across guillemot populations (Fig. 3), with an overall silhouette score of 0.35. The first and largest cluster (n = 51, silhouette score: 0.32) was centred at (0.4 ± 0.6 Z_PFCA_, −0.7 ± 0.6 Z_PFOS_), indicating variable PFCA variability but consistently low PFOS variability. The second cluster (n = 44, silhouette score: 0.37) displayed the opposite pattern, with consistently low PFCA variability but more variable PFOS variability, centred at (−0.9 ± 0.6 Z_PFCA_, 0.3 ± 0.7 Z_PFOS_). The smallest cluster (n = 17, silhouette score: 0.23) showed high variability in both PFAS classes, centred at markedly positive values (1.1 ± 0.8 Z_PFCA_, 1.4 ± 0.9 Z_PFOS_).

**Figure 3.**
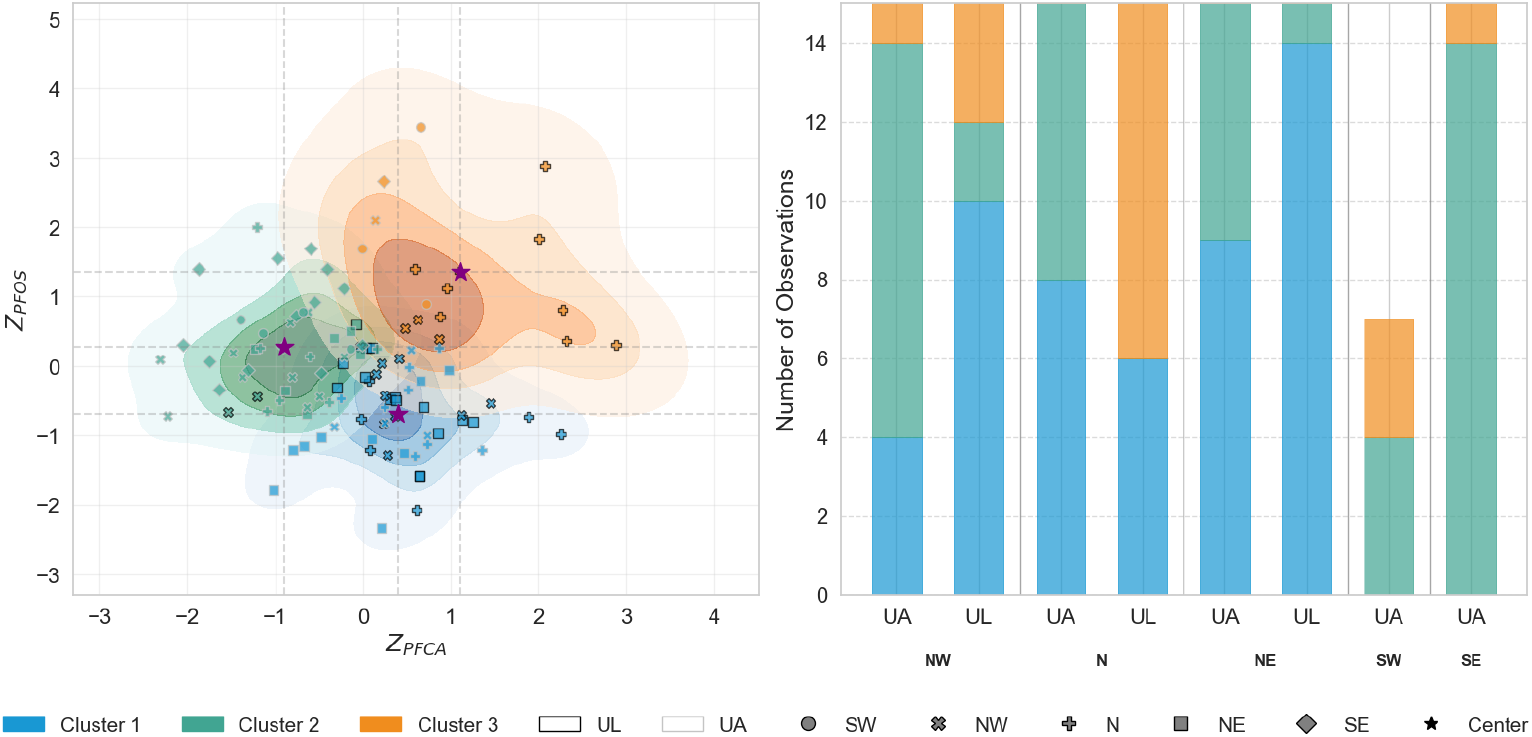
PFAS variability analysis in UA and UL. Left: K-means clustering of standardized PFAS variability (Z-scores). The x-axis (Z_PFCA_) represents variability in long-chain PFCAs (C9-C13), while the y-axis (Z_PFOS_) represents variability in PFOS. The zero point on each axis represents the mean variability, with positive and negative values indicating higher and lower variability than the mean, respectively. Each point represents an individual seabird, with edge colours denoting species (black: UL; white: UA), shapes indicating colony location, and fill colours representing cluster membership. Purple stars mark cluster centroids. Right: Distribution of clusters across colonies, showing the proportion of individuals from each species assigned to each cluster at different colony locations. Colony codes: NW (northwest), N (north), NE (northeast), SW (southwest), SE (southeast).

PFAS variability patterns showed a clear north-south gradient, with Cluster 1 (variable PFCAs, low PFOS variability) dominating northern colonies and Cluster 2 (low PFCA, variable PFOS variability) predominating southern colonies. Species distributions varied by location: at NW colony, UA individuals mainly fell into Cluster 2 while UL individuals primarily grouped in Cluster 1; at colony N, both species shared representation in Cluster 1 but diverged in secondary patterns (UA in Cluster 2, UL in Cluster 3); and at NE colony, both species predominantly occupied Cluster 1 with minimal divergence.

### Isotopic Consistency Across Tissues

Blood cell, δ^13^C values showed a north-south gradient ranging from −22‰ to −19‰, with more negative values in northern colonies and more positive values in southern colonies (Fig. 4). Plasma δ^13^C patterns resembled those of the cell, but with slightly more negative overall values. Blood δ^15^N values ranged from 11‰ to 14‰, and plasma from 11 to 15‰. Both tissues displayed the highest δ^15^N values in the SE colony. Detailed statistical results are in the Supporting Information (Table. S7).

**Figure 4.**
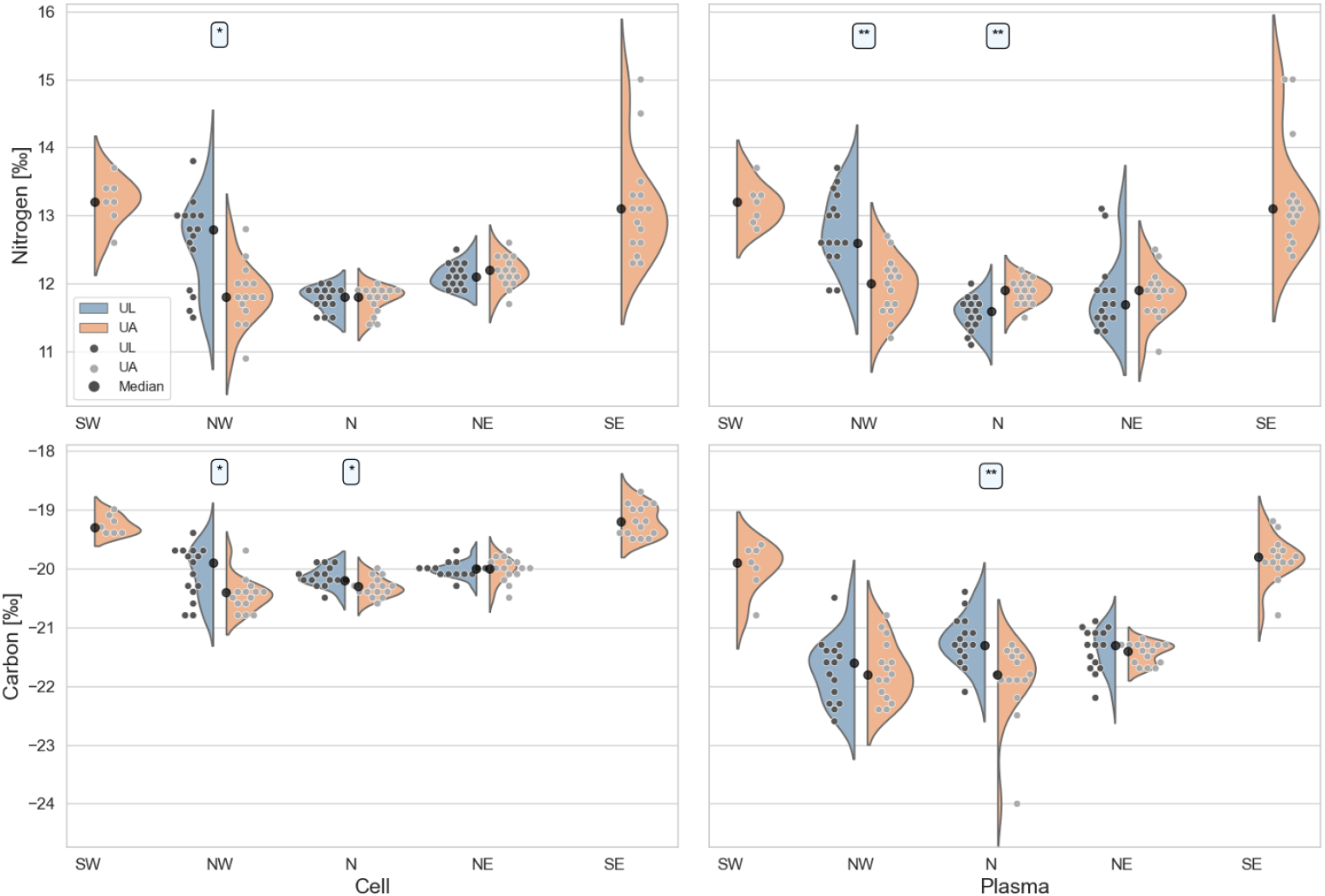
Split Violin Plots of Isotopic Distributions of δ^13^C (Upper Row) and δ^15^N (Lower Row) Across Red Blood Cells (Left Column) and Plasma (Right Column). Each subplot presents the isotopic distribution of δ^13^C (upper) and δ^15^N (lower) in RBCs (left) and plasma (right) in UA and UL across colonies. The split violin plots illustrate the density and distribution of isotopic values, with comparisons made between sympatric colonies. Statistical significance of interspecies comparisons is annotated as follows: *p* < 0.05, ‘*’; *p* < 0.001, ‘**’; *p* < 0.0001, ‘***’.

To assess foraging consistency during the breeding season, we examined isotopic relationships between plasma and red blood cells. Both δ^13^C (*r* = 0.65, *p* < 0.001) and δ^15^N (*r* = 0.70, *p* < 0.001) values showed significant positive correlations between tissues, indicating temporal consistency in foraging patterns. To quantify this consistency across both isotopes simultaneously, we applied PCA to the dual-tissue isotope data, capturing the shared variability between plasma and red blood cell measurements. We then transformed PC1 scores into standardized Z-scores, denoted as δ^15^N_consist_ and δ^13^C_consist_. Stable isotope consistency patterns varied across species and colonies. For nitrogen (δ^15^N_consist_), UA breeding at southernmost colonies showed predominantly positive Z-scores (median SE: 1.3, SW: 1.4), indicating more positive δ^15^N values consistently across both tissues. In contrast, northernmost colonies of both species exhibited negative Z-scores (ranging from −0.9 to −0.2), reflecting more negative δ^15^N values across tissues. Carbon isotope consistency (δ^13^C_consist_) showed similar geographic patterns, with UA from southernmost colonies displaying positive Z-scores (median SE: 2.0, SW: 1.6) indicating consistently more positive δ^13^C values, while northern populations of both species showed negative Z-scores (ranging from −0.9 to −0.1), reflecting consistently more negative δ^13^C values across plasma and cell tissues. Both isotope systems demonstrated consistent north-south gradients, with UL showing relatively similar patterns to northernmost UA.

### Bivariate Segmented Regression

The bivariate segmented regression analyses revealed that PFCA and PFOS variability patterns respond differently to oceanographic thresholds (Table 2). Statistical breakpoints (δ^13^C_consist_ = 0.19, δ^15^N_consist_ = 0.00) delineated three distinct foraging habitats in isotopic space: Arctic-influenced, niche partitioning, and Atlantic-influenced waters (Fig. 5). PFOS variability (Z_PFOS_) showed a clear oceanographic threshold effect, with a statistically significant shift at δ^13^C_consist_ = 0.19 (*p* < 0.01). Mean Z_PFOS_ scores were negative in Arctic-influenced habitats, indicating constrained variability, but strongly positive in Atlantic-influenced habitats, reflecting more variable patterns. This habitat effect accounted for 66% of the total model influence on PFOS variability. In contrast, PFCA variability (Z_PFCA_) showed a more gradual transition across the oceanographic gradient, with non-significant threshold effects. Mean Z_PFCA_ scores were positive in Arctic waters but became increasingly negative in Atlantic waters. While not statistically significant, the δ^15^N_consist_ threshold accounting for 55% of the total model effect. The niche partitioning segment contained predominantly UL individuals from the NW colony, exhibiting negative values for both Z_PFCA_ and Z_PFOS_. These contrasting patterns demonstrate that PFCA and PFOS respond to different ecological drivers: PFCA variability is more influenced by trophic position (δ^15^N_consist_), while PFOS variability is predominantly controlled by habitat factors (δ^13^C_consist_). Detailed statistical results are in the Supporting Information (Table. S8-9).

**Table 2.**
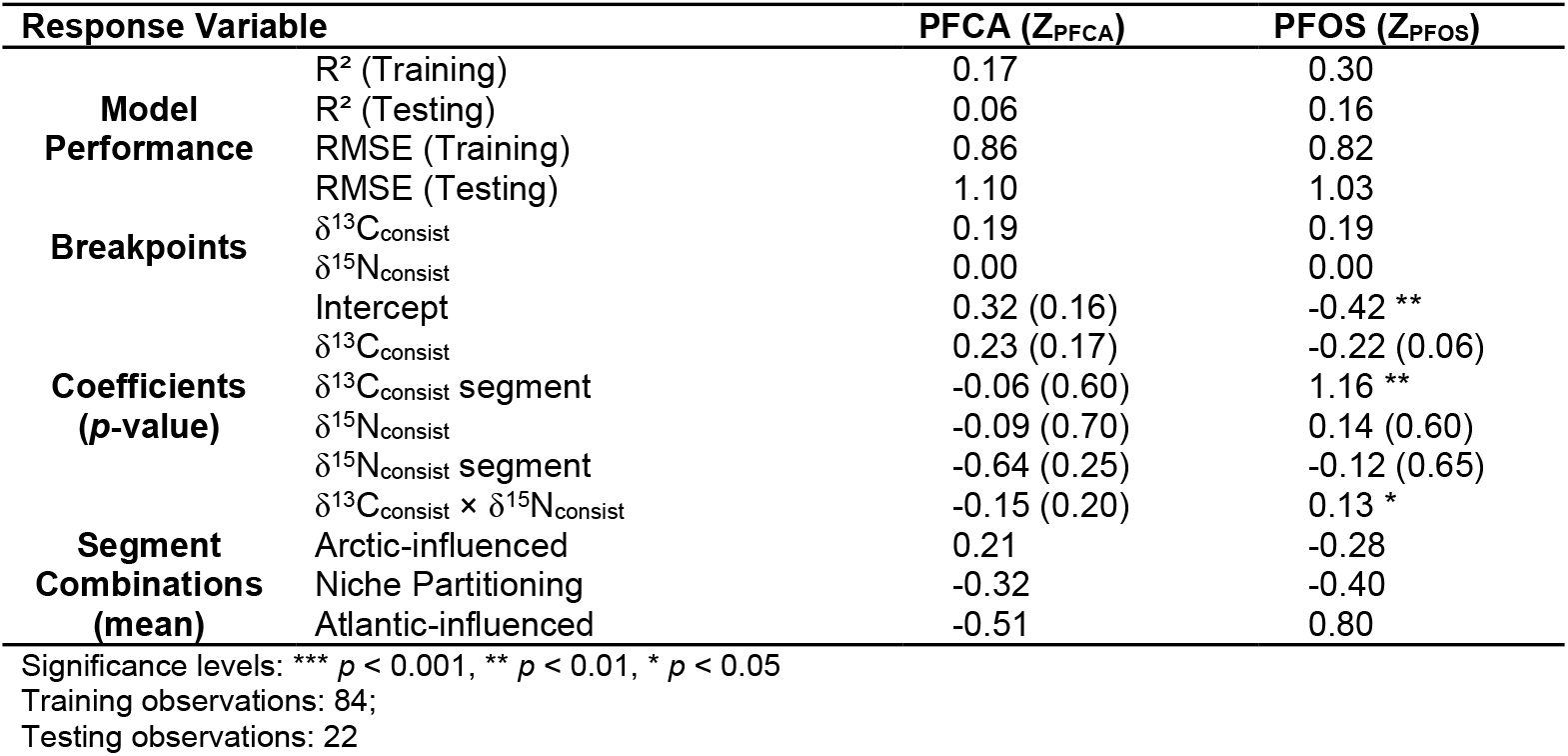
Bivariate Segmented Regression Analysis of Isotopic Indicators’ Influence on PFAS Variability Patterns.

**Figure 5.**
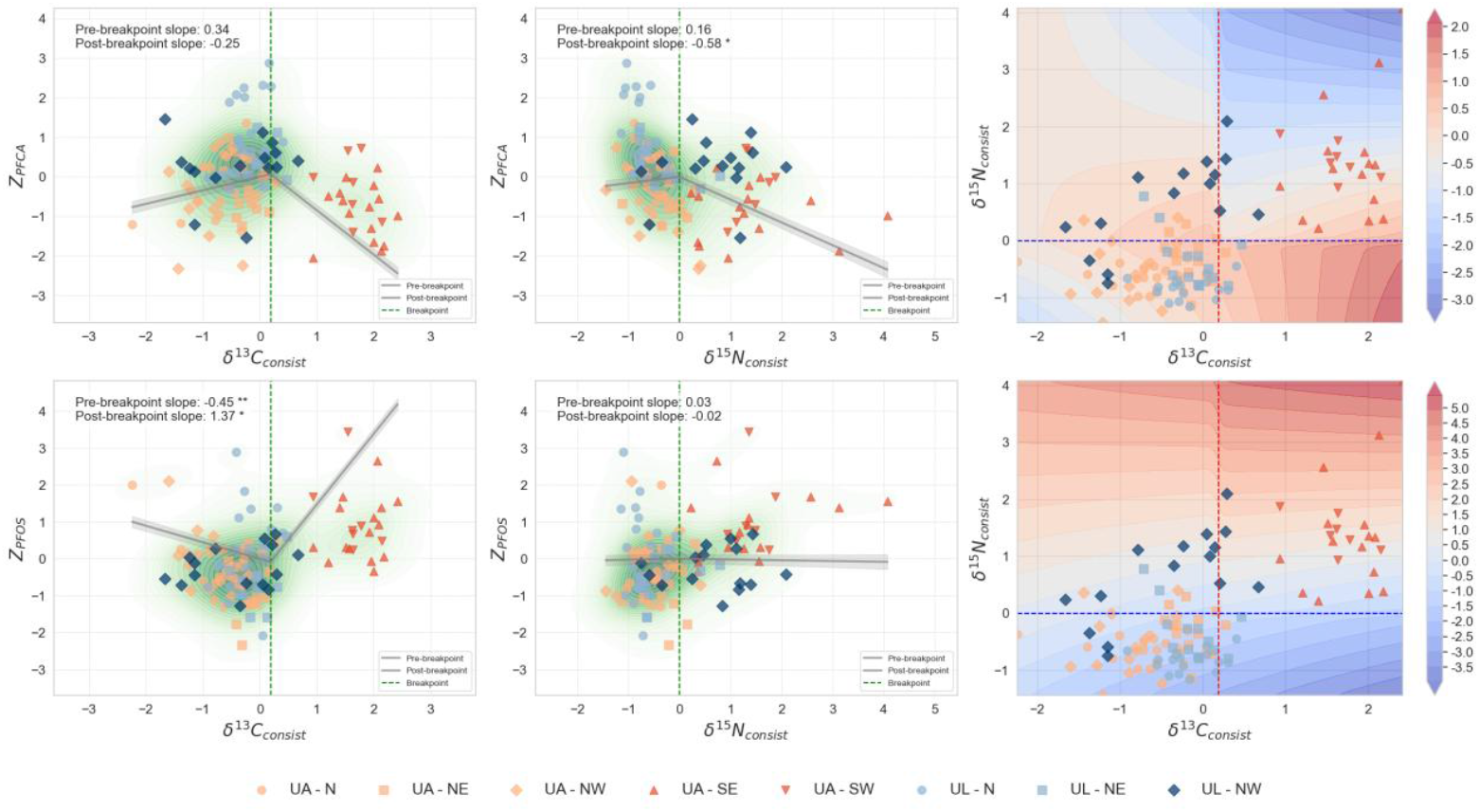
Discrete segmented regression and interaction effects. Segmented regression plots showing the relationship between PFAS variability s and foraging consistency for two models, with regression lines indicating breakpoints (dashed green lines) and pre/post-breakpoint effects (numbers in boxes, asterisks indicate significance levels: \**p* < 0.05, \*\**p* < 0.01, \*\*\**p* < 0.001). Contour plots displaying the interaction between predictors, with breakpoints shown as dashed red (vertical, δ^13^C_consist_ = 0.19) and blue (horizontal, δ^15^N_consist_ = 0.00) lines. The resulting quadrants represent distinct foraging habitats: Arctic-influenced (lower left, δ^13^C_consist_ < 0.19 and δ^15^N_consist_ < 0.00), niche partitioning from Arctic-influenced (upper left, δ^13^C_consist_ < 0.19 and δ^15^N_consist_ > 0.00), and Atlantic-influenced (upper right, δ^13^C_consist_ > 0.19 and δ^15^N_consist_ > 0.00). Colour gradients represent response values from −2 to 2 in 0.25 increments. Species are indicated by colour (UL: blue, UA: orange) and colonies by markers.

## Discussion

PFAS bioaccumulation patterns in marine predators reflect the dynamic integration between environmental exposure pathways and trophic transfer processes.^5, 6^ Whereas previous studies have treated variability as statistical noise, our approach recognises that variability as ecologically informative, revealing aspects of contaminant pathways that mean-based analyses may miss.^5, 6, 11^ Our integrated dual-isotope framework (δ^13^C_consist_ and δ^15^N_consist_ scores) allows us to interpret PFAS variability as an ecological signal, showing how foraging patterns influence contamination dynamics. Despite inherent methodological limitations as discussed later, this approach offers insights into the ecological drivers of contaminant exposure.^38, 67^ Our variability-based framework demonstrates broader applicability beyond PFAS, potentially revealing ecological processes governing other persistent contaminants in marine food webs where spatial heterogeneity and foraging specialisation create measurable variability patterns. For instance, the segmented regression analysis (Table 2, Fig. 5) identified significant isotopic thresholds aligning with Arctic-Atlantic oceanographic regimes. These thresholds reveal how variability patterns shift across these oceanographic gradients even when mean concentrations might not show clear transitions. Similarly, cluster analysis (Fig. 3) identified habitat-specific variability differences between sympatric species with pronounced niche separation at the NW colony that traditional approaches would mask. These findings demonstrate how variability patterns uncover ecological drivers, such as the different influences of trophic position on long-chain PFCAs and habitat use on PFOS, thus contributing to our understanding of contaminant dynamics in rapidly changing marine ecosystems.

### PFAS Variability as Structured Ecological Signal

Our hypotheses predicted that (1) PFAS exposure variability contains structured ecological signals reflecting foraging strategies and oceanographic influences, and (2) UL and UA would exhibit distinct variability profiles, with UL showing more constrained patterns and UA exhibiting broader variability. The results provide support for structured ecological signals while revealing that species differences are compound- and habitat-dependent rather than universally expressed. K-means clustering revealed three distinct variability patterns with clear geographic organization (Fig. 3), while significant threshold effects at Arctic-Atlantic transitions (δ^13^C_consist_ = 0.19, δ^15^N_consist_ = 0.00) mark ecological boundaries. The temporal consistency in foraging behaviour during breeding (plasma-cell isotope correlations: *r* = 0.65 - 0.70) indicates that individuals maintain relatively stable foraging patterns over tissue integration periods, enabling confident interpretation of relationships between current foraging indicators and PFAS variability patterns despite temporal mismatches between isotopic integration (weeks - months) and PFAS accumulation (years).

As expected from previously published telemetry data^68^, these geographic patterns demonstrate that PFAS variability reflects structured ecological signals rather than statistical noise. Both guillemot species breeding in northern Iceland exhibited high site fidelity by foraging in cold, less saline Arctic waters, reflected in more negative δ^13^C_consist_ and δ^15^N_consist_ scores (Fig. 4) and primarily associated with constrained PFOS variability (Fig. 3). In contrast, southern Icelandic UA foraged within warmer, more saline Atlantic waters (evidenced by more positive isotopic scores), demonstrating constrained PFCA variability yet more variable PFOS patterns. This compound-specific response validates distinct bioaccumulation mechanisms operating within the sister species.

While these broad-scale patterns demonstrate how water masses influence PFAS variability, our detailed analysis identified fine-scale habitat-specific behavioural adaptations that generated notable differences from expected patterns. At the NW colony, the variability patterns of both UA and UL distributed across three clusters, with UL showing predominantly constrained PFOS variability whereas UA showed constrained PFCA variability (Fig. 3), reflecting their differences in foraging strategies. As shown in our tracking data^68^, oceanographic heterogeneity at the interface between the warm Irminger Current and cold East Greenland Current enabled distinct foraging opportunities where UL foraged more frequently in glacial fjords and along the marginal ice zone compared to UA^58^, while they also showed substantial overlaps in their use of other habitat features such as water depth. This pattern suggests that species-specific foraging behaviours may influence compound-specific variability patterns. While the mechanisms remain to be fully elucidated, the species-compound coupling demonstrates that environmental heterogeneity enables the expression of different variability patterns rather than simply creating broad variability differences. At the N colony, while both species shared more similar foraging strategies and habitat use^68^, UA showed more defined PFAS variability patterns compared to UL (Fig. 3). Within this colony, both UA and UL were associated with constrained PFOS variability. However, a considerable portion of UA showed constrained PFCA variability, whereas more than half of the observed UL showed less defined variability (Fig. 3). Diel vertical migration of marine prey provides the ecological foundation for how depth- and time-specific foraging may expose the two species to different prey assemblages and contamination sources.^69, 70^ The documented differences between species, UL performing nocturnal foraging and shallower diving while UA dived to deeper depths throughout the day^68^, position them to encounter different components of the vertically migrating prey community. This temporal-depth decoupling potentially creates the observed PFAS variability patterns, particularly explaining UL’s less defined variability. At the NE colony, the variability patterns were the most uniform, as both UA and UL are predominantly associated with constrained PFOS variability. Within NE, no niche partitioning was observed between species.^58^ This ecological convergence reflects both species accessing similar prey resources within Arctic food web structures, and create homogeneous exposure environments.

In contrast, the SW and SE colonies demonstrate how trophic magnification processes create compound-specific uniform variability patterns despite isotopic diversity. UA at these colonies exhibited the highest δ^15^N values across both tissues (Fig. 4), indicating foraging at relatively high trophic levels within Atlantic-influenced waters. Correspondingly, the highest PFOS concentrations were observed within these colonies (Fig. 2), indicating enhanced biomagnification processes during high trophic-level foraging.^6, 8^ This enhanced biomagnification creates compound-specific responses through different mechanisms. For PFOS, the elevated concentrations enable greater detection of inter-individual variability, resulting in more variable PFOS patterns. For PFCAs, however, the intense biomagnification results in competitive protein-binding where PFOS consistently outcompetes PFCAs for limited binding sites regardless of prey diversity consumed.^71, 72^ This competitive dominance suppresses PFCA accumulation and constrains PFCA variability, creating uniform PFCA patterns. The result is compound-specific variability patterns where bioaccumulation intensity differentially affects PFOS and PFCA detectability through distinct mechanisms, concentration-enhanced detection for PFOS versus competition-mediated suppression for PFCAs.

On the other hand, birds at Arctic-influenced colonies foraged on lower trophic sources with reduced biomagnification compared to Atlantic systems, creating contrasting effects on variability patterns. For PFOS, lower concentrations from reduced biomagnification constrain detectable inter-individual variability, creating uniform patterns. For PFCAs, the absence of intense biomagnification prevents competitive protein-binding, making PFCA variability more detectable rather than being biochemically suppressed. Arctic systems thus preserve ecological signals as primary drivers of PFCA variability while PFOS variability becomes constrained by both ecological similarity and concentration-dependent detection. This represents ecological signal preservation rather than the biochemical masking observed in Atlantic systems.

### Methodological Considerations

Our approach of using temporal consistency in isotopic values (δ^13^C_consist_ scores and δ^15^N_consist_ scores) for foraging behaviour provides a framework for interpreting PFAS variability patterns across oceanographic gradients. While these consistency scores offer insights into foraging habits during the breeding season, several methodological limitations should be considered when interpreting the results. One consideration is the temporal mismatch between isotope turnover rates (weeks to months) and PFAS persistence (years in some cases). While the isotopes analysed in the two blood fractions reflect recent foraging behaviour, part of the detected PFAS likely accumulated before the breeding season. This temporal integration is particularly relevant for legacy compounds like PFOS, which has a half-life of several years in seabirds^73^. Additionally, isotopic baselines can vary substantially across oceanographic regions, seasons, and even small spatial scales.^74, 75^ Without direct measurements of baseline isotopic values across our study area, we cannot definitively attribute differences in isotopic values to trophic or habitat factors versus baseline shifts. For example, δ^15^N values may reflect differences in baseline nitrogen sources rather than actual trophic positions.^76^ Physiological factors unrelated to diet can also influence both isotopic values and PFAS accumulation. Individual variation in metabolic rates, growth, reproductive status, and detoxification capacity may affect both isotopic incorporation and PFAS retention independently, potentially creating correlations that do not reflect ecological processes. Finally, our correlative approach cannot establish causation between isotopic patterns and PFAS variability. While our 2018 breeding season data capture apparent habitat associations, these relationships require experimental validation to confirm the mechanistic links between foraging ecology and PFAS accumulation patterns. Nevertheless, several methodological advances could address these limitations. Isotope analysis of tissues formed before the breeding season and inclusion of more sophisticated analyses such as compound specific isotope analyses that can provide more detailed insight into foraging and physiology compared to bulk stable isotope analyses. ^25, 77–79, 80–82^ Furthermore, baseline corrections across different oceanographic regions could improve the comparability of isotopic measurements between Arctic and Atlantic waters.^83^

## Conclusion and Implications

Our integrated analysis highlights how PFAS exposure variability in marine predators is structured by a combination of oceanographic features, foraging ecology, and compound-specific behaviour. Spatial habitat factors predominantly shape PFOS variability, with pronounced transitions at Arctic-Atlantic boundaries, while long-chain PFCA variation aligns with trophic indicators, reflecting foraging position within food webs. These patterns point to the influence of both large-scale oceanographic gradients and colony-level foraging behaviours in generating local variation in exposure. The distinct PFOS and PFCA responses reflect their unique physicochemical properties and biological interactions. Our variability-focused approaches reveal ecological dimensions of contaminant exposure that mean-based methods typically miss, though temporal mismatches between isotopic integration and PFAS persistence require careful interpretation. As climate change shifts ocean conditions, the oceanographic drivers of PFAS variability we identify will likely alter contaminant risks for marine predators. As demonstrated in this study, this framework offers potential for tracking contaminant pathways and for informing conservation strategies in rapidly changing polar and subpolar marine ecosystems.

## CRediT authorship contribution statement

S.R.: Writing - Review & Editing, Writing - Original Draft, Visualization, Validation, Software, Methodology, Formal Analysis, Data Curation, Conceptualization.

R.E. Conceptualization, Funding Acquisition.

V.D. Writing - Review & Editing, Validation, Methodology.

R.N. Writing - Review & Editing, Funding Acquisition, Conceptualization.

L.T.: Writing - Review & Editing, Funding Acquisition, Formal Analysis, Conceptualization.

## Acknowledgments

This paper is an output of Project LOMVIA (UKRI/NERC Grant No. NE/R012660/1) and Project EISPAC (NERC Grant No. NE/R012857/1), part of the Changing Arctic Ocean Programme. Additional funding was provided by BMBF (Grant No. 03V01459 for LOMVIA, Grant No. 03F0809A for EISPAC) in Germany, with contributions from the Max Planck Society. A permit was provided by the Icelandic Institute of Natural History and animal ethics was overseen by BAS AWERB.

## Appendix A. Supplementary Data

