## supplementary information for "Arctic-Atlantic gradient shaped PFAS exposure variability in sympatric guillemot species off Iceland"

### 1 Methods and Materials

#### 1.1 Sample Collection

Following centrifugation, plasma aliquots were transferred into borosilicate-glass Petri dishes that had been rinsed with deionized water and pre-combusted at 500°C for 30 minutes to minimize PFAS sorption.^1^ Samples were covered with pre-combusted glass lids and placed in a polyethylene vacuum desiccator for 4-6 days of ambient temperature drying to exclude airborne contaminants. Once dry, dishes were sealed with Parafilm, double-bagged in PFAS-free polyethylene zip-bags, and stored at room temperature until extraction.

#### 1.2 Target Compounds & Standards

**Table S1.** PFAS analytes. Chemical information of target per- and polyfluoroalkyl substances (PFAS) analysed in this study. The compounds are categorized into three major classes: carboxylic acids (C4-C13), sulfonic acids (C4, C6, C8, & C10), and ether (HFPO-DA). All analytical standards were obtained from Wellington Laboratories with certified purity >98%. The PFC-MXA mixture was prepared at 2.0 µg/mL ± 5%, while the HFPO-DA individual standard was prepared at 50 ± 2.5 µg/mL.

| **Class** | **Acronym** | **Chemical Name** | **CAS No.** | **Standard Information** |
| --- | --- | --- | --- | --- |
| Carboxylic Acids | PFPeA | perfluoro*-n-*pentanoic acid | 2706-90-3 | PFC-MXA mixture, |
|  | PFHxA | perfluoro*-n-*hexanoic acid | 307-24-4 | Wellington Laboratories, |
|  | PFHpA | perfluoro*-n-*heptanoic acid | 375-85-9 | 2.0 µg/mL ± 5%, |
|  | PFOA | perfluoro*-n-*octanoic acid | 335-67-1 | >98% |
|  | PFNA | perfluoro*-n-*nonanoic acid | 375-95-1 |  |
|  | PFDA | perfluoro*-n-*decanoic acid | 335-76-2 |  |
|  | PFUnDA | perfluoro*-n-*undecanoic acid | 2058-94-8 |  |
|  | PFDoDA | perfluoro*-n-*dodecanoic acid | 307-55-1 |  |
|  | PFTrDA | perfluoro*-n-*tridecanoic acid | 72629-94-8 |  |
| Sulfonic Acids | PFBS | potassium perfluoro*-n-*butanesulfonate | 375-73-5 |  |
|  | PFHxS | sodium perfluoro-*n-*hexanesulfonate | 355-46-4 |  |
|  | PFOS | sodium perfluoro*-n-*octanesulfonate | 1763-23-1 |  |
|  | PFDS | sodium perfluoro*-n-*decanesulfonate | 335-77-3 |  |
| Ether | HFPO-DA | 2,3,3,3-tetrafluoro-2-(1,1,2,2,3,3,3-heptafluoropropoxy)propanoic acid | 13252-13-6 | Individual standard, Wellington Laboratories, |
|  |  |  |  | (50 ± 2.5) µg/mL, |
|  |  |  |  | >98% |

**Table S2.** List of solvents and reagents used in the analytical procedure. All chemicals were of high analytical grade suitable for LC-MS analysis or equivalent high-purity grade. Ultrapure water was produced in-house using a Milli-Q water purification system. The reagent grade, purity specifications, and suppliers are provided to ensure reproducibility of the analytical method.

| **Chemical** | **Grade/Purity** | **Supplier** |
| --- | --- | --- |
| Methanol | LC-MS grade (LiChrosolv) | Merck |
| Ultrapure water | 18.2 MΩ·cm at 25 °C | Milli-Q Integral 5, Merck |
| Acetic acid | ≥99.8%, LC-MS grade | Honeywell Fluka |
| Ammonium acetate | LC-MS grade | Honeywell Fluka |
| Ammonia solution | 25%, Suprapur | Merck |

#### 1.3 Sample Preparation

PFAS extraction from plasma samples was performed using a modified quaternary ammonium salt-based ion-pairing method.^2, 3^ Prior to extraction, plasma samples were fortified with mass‑labelled internal standards and buffered with sodium carbonate (1 mL, 0.5 M, pH 10.0). Tetrabutylammonium hydrogen sulphate (2 mL, 0.5 M) was added as the ion-pairing agent.

The analytes were extracted three times with ethyl acetate (5 mL per extraction) using ultrasonication (20 min, room temperature) followed by centrifugation (3000 rpm, 15 min, 20°C). The combined organic extracts were concentrated under a gentle nitrogen stream at 40°C and reconstituted in methanol: water (1:1 v/v, 250 µL). Final extracts were filtered through a 0.2 µm polypropylene membrane filter prior to analysis.

#### 1.4 Instrumental Analysis

**Table S3.** Detailed instrumental parameters for the HPLC-MS/MS analysis of PFAS . The chromatographic separation was performed using an Agilent 1100 HPLC system coupled to an AB Sciex API 4000 triple quadrupole mass spectrometer. The method utilized a reverse-phase C18 column with gradient elution using ammonium acetate buffer and methanol-based mobile phases. Mass spectrometric detection was performed using electrospray ionization in negative mode with multiple reaction monitoring (MRM).

| **System** | **Parameter** | **Specification** |
| --- | --- | --- |
| **HPLC** | System | HP 1100, Agilent Technologies |
|  | Analytical Column | Synergi Fusion-RP C18, 150 × 2 mm, 4 µm, 80 Å (Phenomenex) |
|  | Guard Column | SecurityGuard C18, 4 × 2 mm (Phenomenex) |
|  | Mobile Phase A | 2 mM ammonium acetate in water |
|  | Mobile Phase B | 0.05% acetic acid in methanol |
|  | Flow Rate | 0.2 mL/min |
|  | Injection Volume | 10 µL |
|  | Column Temperature | 30 °C |
|  | Run Rime | 30 min |
| **MS/MS** | System | API 4000 triple quadrupole, AB Sciex |
|  | Source | Turbo V ESI, negative mode |
|  | Ion Spray Voltage | -4500 V |
|  | Source Temperature | 400 °C |
|  | Gases (N_2_) | Nebulizer: 4.2 bar |
|  |  | Heater: 2.8 bar |
|  |  | Curtain: 1.0 bar |
|  |  | Collision: 0.6 bar |
|  | Scan Type | MRM |

**Table S4.** LC-MS/MS parameters for the analysis of per- and polyfluoroalkyl substances (PFAS) and their corresponding internal standards. The table presents molecular formulas of detected ions, quantifier and qualifier transitions (m/z), and matched internal standards for each target analyte. The compounds are categorized into native PFCAs (C4-C13), PFSAs (C4, C6, C8 & C10), HFPO-DA, and isotope‑labelled internal standards. Quantifier ion transitions (marked with asterisk) were used for quantitation, while qualifier transitions were monitored for confirmation where available.

| Type | Compound | Molecular Formula | Quantifier Transition (m/z) | Qualifier Transition (m/z) | Internal Standard |
| --- | --- | --- | --- | --- | --- |
| Native PFCAs | PFPeA | [C5F9O2]- | 263 > 219* | - | ^13^C_2_-PFHxA |
|  | PFHxA | [C6F11O2]- | 313 > 269* | 313 > 119 | ^13^C_2_-PFHxA |
|  | PFHpA | [C7F13O2]- | 363 > 169* | 363 > 319 | ^13^C_4_-PFOA |
|  | PFOA | [C8F15O2]- | 413 > 369* | 413 > 169 | ^13^C_4_-PFOA |
|  | PFNA | [C9F17O2]- | 463 > 419* | 463 > 219 | ^13^C_5_-PFNA |
|  | PFDA | [C10F19O2]- | 513 > 469* | 513 > 219 | ^13^C_2_-PFDA |
|  | PFUnDA | [C11F21O2]- | 563 > 519* | 563 > 169 | ^13^C_2_-PFUnDA |
|  | PFDoDA | [C12F23O2]- | 613 > 569* | 613 > 169 | ^13^C_2_-PFDoDA |
|  | PFTrDA | [C13F25O2]- | 663 > 619* | 663 > 169 | ^13^C_2_-PFDoDA |
| Native PFSAs | PFBS | [C4F9O3S]- | 299 > 80* | 299 > 99 | ^18^O_2_-PFHxS |
|  | PFHxS | [C6F13O3S]- | 399 > 80* | 399 > 99 | ^18^O_2_-PFHxS |
|  | PFOS | [C8F17O3S]- | 499 > 80* | 499 > 99 | ^13^C_4_-PFOS |
|  | PFDS | [C10F21O3S]- | 599 > 80* | 599 > 99 | ^13^C_4_-PFOS |
| Native Ether | HFPO-DA | [C6F11O3]- | 329 > 285 | - | ^13^C_3‑_HFPO‑DA |
| Internal Standards | ^13^C_2_-PFHxA | [13C2C4F11O2]- | 315 > 270* | 315 > 120 | - |
|  | ^13^C_4_-PFOA | [13C4C4F15O2]- | 417 > 372* | 417 > 169 | - |
|  | ^13^C_8_-PFOA | [13C8F15O2]- | 421 > 376* | 421 > 172 | - |
|  | ^13^C_5_-PFNA | [13C5C4F17O2]- | 468 > 423* | 468 > 223 | - |
|  | ^18^O_2_-PFHxS | [C6F13O2OS]- | 403 > 84* | 403 > 103 | - |
|  | ^13^C_4_-PFOS | [13C4C4F17O3S]- | 503 > 80* | 503 > 99 | - |
|  | ^13^C_3‑_HFPO‑DA | [13C3C3F11O3]- | 332 > 287* | 332 > 169 | - |
| *Quantifier ion transition | | | | | |

## 1.5 QA/QC

Background PFAS levels were evaluated through procedural blanks analysed with each batch of 10 samples. The Limit of Blank (LoB) was calculated as^4^:

$$LoB=mean(blank)+1.645(SDblank)$$

Sample concentrations above LoB were considered detectable and were blank corrected by subtracting the mean blank values. Values below LoB were reported as not detected. Recovery rates for all target compounds and their mass-labelled internal standards are summarized in Table A5.

**Table S5.** Mean percent recoveries and standard deviations (SD) for isotope-labelled per- and polyfluoroalkyl substances (PFAS) internal standards.

| **Standard** | **Mean** | **SD** |
| --- | --- | --- |
| ^13^C_2_-PFHxA | 78% | 10% |
| ^13^C_4_-PFOA | 88% | 11% |
| ^13^C_5_-PFNA | 87% | 11% |
| ^13^C_2_-PFDA | 86% | 17% |
| ^13^C_2_-PFUnDA | 87% | 22% |
| ^13^C_2_-PFDoDA | 102% | 17% |
| ^18^O_2_-PFHxS | 93% | 12% |
| ^13^C_4_-PFOS | 102% | 14% |
| ^13^C_4_-PFDS | 94% | 13% |
| ^13^C_3‑_HFPO‑DA | 131% | 80% |

Calibration curves showed good linearity (*R*² > 0.995) over the concentration range of 0.0 - 100 pg/µL. Method detection limits (MDL) were determined based on signal-to-noise ratio calculations following the guidelines outlined in Agilent's technical report on mass spectrometry detection limits.^5^ The calculated MDL, LoB and mean blank values for each target compound are detailed in Table A6.

**Table S6.** Method performance parameters for target PFAS

| **Class** | **Acronym** | **MDL [ng/mL]** | **LoB [ng/mL]** | **Blanks [ng/mL]** |
| --- | --- | --- | --- | --- |
| Carboxylic | PFPeA | 0.06 | 0.23 | 0.22 |
| Acids | PFHxA | 0.06 | 0.33 | 0.3 |
|  | PFHpA | 0.06 | < 0.06 | < 0.06 |
|  | PFOA | 0.06 | 0.11 | 0.13 |
|  | PFNA | 0.06 | < 0.06 | < 0.06 |
|  | PFDA | 0.07 | < 0.07 | < 0.07 |
|  | PFUnDA | 0.05 | < 0.05 | < 0.05 |
|  | PFDoDA | 0.05 | < 0.05 | < 0.05 |
|  | PFTrDA | 0.06 | < 0.06 | < 0.06 |
| Sulfonic | PFBS | 0.06 | < 0.06 | < 0.06 |
| Acids | PFHxS | 0.06 | < 0.06 | < 0.06 |
|  | PFOS | 0.05 | 0.34 | 0.28 |
|  | PFDS | 0.06 | < 0.06 | < 0.06 |
| Ether | HFPO-DA | 0.06 | < 0.06 | < 0.06 |

### 2 Stats Results

#### 2.1 PFAS Analysis

##### 2.1.1 PCA

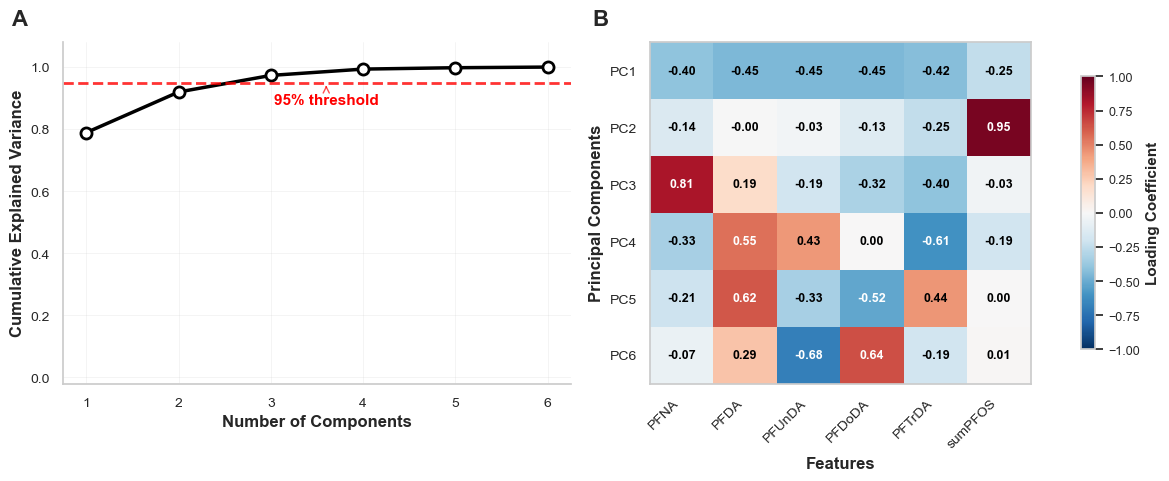

**Figure S1.** Principal Component Analysis (PCA) visualization of the PFAS dataset. (A) Explained variance ratio across principal components, showing the proportion of variance captured by each component. (B) Heatmap depicting feature contributions to the first six principal components, where colour intensity represents loading strength and direction.

#### 2.2 Stable Isotope Analysis

##### 2.2.1 Descriptives of SIR

**Table S7.** Statistical summary of stable isotope values (δ¹⁵N and δ¹³C) in cell and plasma samples from UA (*n* = 67) and UL (*n* = 45) groups.

| **Species** | **Colony** | **δ^15^N_cell_** | | **δ^13^C_cell_** | | **δ^15^N_plasma_** | | **δ^13^C_plasma_** | |
| --- | --- | --- | --- | --- | --- | --- | --- | --- | --- |
|  |  | **Median** | **IQR** | **Median** | **IQR** | **Median** | **IQR** | **Median** | **IQR** |
| UA | N | 11.8 | 0.3 | -20.3 | 0.1 | 11.9 | 0.2 | -21.8 | 0.4 |
| UA | NE | 12.2 | 0.3 | -20.0 | 0.2 | 11.9 | 0.4 | -21.4 | 0.3 |
| UA | NW | 11.8 | 0.4 | -20.4 | 0.2 | 12.0 | 0.6 | -21.8 | 0.7 |
| UA | SE | 13.1 | 0.7 | -19.2 | 0.4 | 13.1 | 0.4 | -19.8 | 0.2 |
| UA | SW | 13.2 | 0.3 | -19.3 | 0.2 | 13.2 | 0.4 | -19.9 | 0.4 |
| UL | N | 11.8 | 0.2 | -20.2 | 0.1 | 11.6 | 0.3 | -21.3 | 0.4 |
| UL | NE | 12.1 | 0.2 | -20.0 | 0.1 | 11.7 | 0.4 | -21.3 | 0.5 |
| UL | NW | 12.8 | 0.8 | -19.9 | 0.7 | 12.6 | 0.7 | -21.6 | 0.8 |

##### 2.2.2 Correlation Analysis

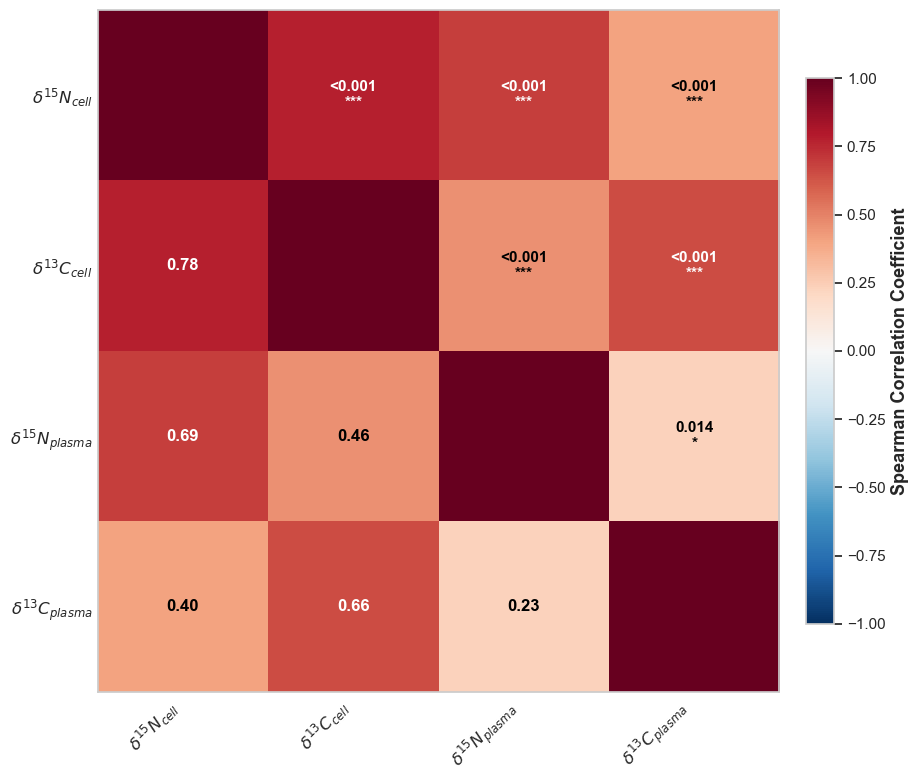

**Figure S2.** Spearman Correlation Heatmap of stable isotope values across tissue types correlation matrix illustrating the relationships between carbon (δ¹³C) and nitrogen (δ¹⁵N) isotope values in cell and plasma tissues. Colour intensity represents correlation strength, with red indicating positive correlations and blue indicating negative correlations. Asterisks denote statistical significance levels: *** *p* < 0.001, ** *p* < 0.01, * *p* < 0.05, ns *p* ≥ 0.05. Values within each cell represent the Spearman correlation coefficient.

##### 2.2.3 PCA

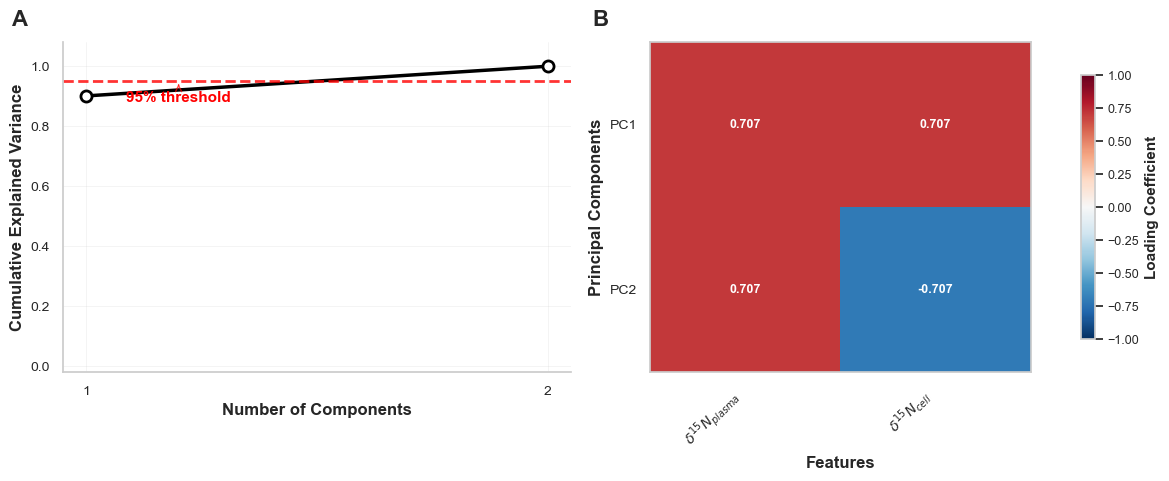

**Figure S3.** Principal Component Analysis of δ^15^N in plasma and cells. **(A)** Scree plot showing cumulative explained variance by principal components. The red dashed line indicates the 95% variance threshold. **(B)** Feature contributions (loadings) to principal components PC1 and PC2. Loading coefficients range from -1 to +1, with red indicating positive loadings and blue indicating negative loadings. PC1 explains 90.1% of the total variance, while PC2 explains the remaining 9.9%.

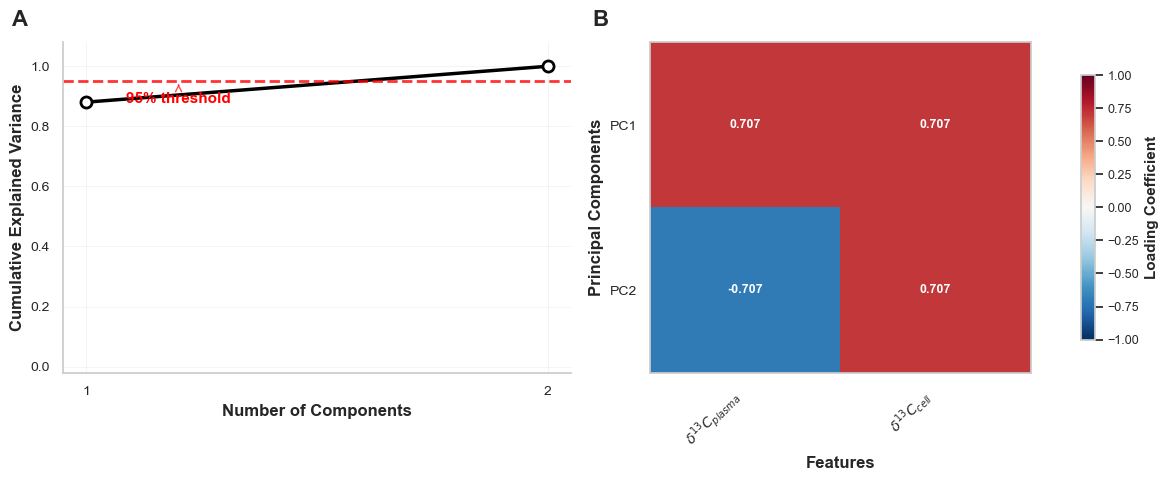

**Figure S4.** Principal Component Analysis of δ^13^C in plasma and cells. **(A)** Scree plot showing cumulative explained variance by principal components. The red dashed line indicates the 95% variance threshold. **(B)** Feature contributions (loadings) to principal components PC1 and PC2. Loading coefficients range from -1 to +1, with red indicating positive loadings and blue indicating negative loadings. PC1 explains 88.1% of the total variance, while PC2 explains the remaining 11.9%.

#### 2.3 Bivariate Segmented Regression Analysis

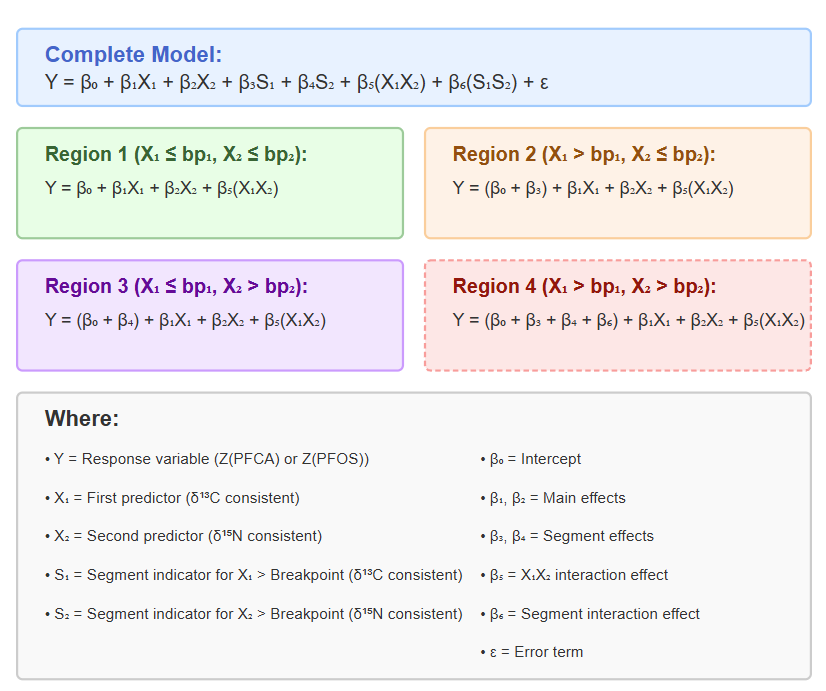

**Figure S5.** Segmented regression model with dual breakpoints for isotope values (δ¹³C and δ¹⁵N). Due to small sample size, Region 4 was excluded from the analysis; the model was implemented using only Regions 1, 2, and 3 combined.

**Table S8.** Bivariate Segmented Regression Results (PFCA)

| **Component** | **Parameter** | **Value** | **SE** | ***t*** | ***p*** | **Training** | **Testing** | **% Effect** | **n** | **Mean** | **SD** | **Min** | **Max** |
| --- | --- | --- | --- | --- | --- | --- | --- | --- | --- | --- | --- | --- | --- |
| **Model Info** | Response Variable | Z_PFCA_ | - | - | - | - | - | - | - | - | - | - | - |
|  | Predictor 1 | δ^13^C_consist_ | - | - | - | - | - | - | - | - | - | - | - |
|  | Predictor 2 | δ^15^N_consist_ | - | - | - | - | - | - | - | - | - | - | - |
|  | Training Observations | 89 | - | - | - | - | - | - | - | - | - | - | - |
|  | Testing Observations | 23 | - | - | - | - | - | - | - | - | - | - | - |
| **Performance** | *R*² | - | - | - | - | 0.19 | 0.03 | - | - | - | - | - | - |
|  | MSE | - | - | - | - | 0.78 | 1.12 | - | - | - | - | - | - |
|  | RMSE | - | - | - | - | 0.88 | 1.06 | - | - | - | - | - | - |
| **Breakpoints** | δ^13^C_consist_ | 0.19 | - | - | - | - | - | - | - | - | - | - | - |
|  | δ^13^C_consist_ Below | 80 (71.4%) | - | - | - | - | - | - | - | - | - | - | - |
|  | δ^13^C_consist_ Above | 32 (28.6%) | - | - | - | - | - | - | - | - | - | - | - |
|  | δ^15^N_consist_ | 0 | - | - | - | - | - | - | - | - | - | - | - |
|  | δ^15^N_consist_ Below | 71 (63.4%) | - | - | - | - | - | - | - | - | - | - | - |
|  | δ^15^N_consist_ Above | 41 (36.6%) | - | - | - | - | - | - | - | - | - | - | - |
| **Coefficients** | Intercept | 0.54 | - | - | - | - | - | - | - | - | - | - | - |
|  | δ^13^C_consist_ | 0.34 | 0 | 1.96 | 0.05 | - | - | 13.18 | - | - | - | - | - |
|  | δ^13^C_consist_ segment | 0.27 | 1 | 0.5 | 0.62 | - | - | 10.5 | - | - | - | - | - |
|  | δ^15^N_consist_ | 0.16 | 0 | 0.98 | 0.33 | - | - | 6.11 | - | - | - | - | - |
|  | δ^15^N_consist_ segment | -0.74 | 0 | -2.4 | **0.02*** | - | - | 28.32 | - | - | - | - | - |
|  | δ^13^C_consist_ × δ^15^N_consist_ interaction | -0.22 | 0 | -2.4 | **0.02*** | - | - | 8.44 | - | - | - | - | - |
|  | Segment interaction | -0.87 | 1 | -1.2 | 0.23 | - | - | 33.45 | - | - | - | - | - |
| **Segment Combinations** | Low δ^13^C_consist_, Low δ^15^N_consist_ | - | - | - | - | - | - | - | 65 | 0.21 | 1 | -1.5 | 2.9 |
|  | Low δ^13^C_consist_, High δ^15^N_consist_ | - | - | - | - | - | - | - | 15 | -0.32 | 1 | -2.3 | 1.5 |
|  | High δ^13^C_consist_, Low δ^15^N_consist_ | - | - | - | - | - | - | - | 6 | 0.78 | 1 | -0.1 | 2.3 |
|  | High δ^13^C_consist_, High δ^15^N_consist_ | - | - | - | - | - | - | - | 26 | -0.51 | 1 | -2.1 | 0.9 |
| **Note:** Bold *p*-values indicate statistical significance (*p* < 0.05). Asterisks (*) denote significant terms. The model shows poor predictive performance (Testing *R*² = 0.03). δ^15^N_consist_ segment and the δ^13^C_consist_ × δ^15^N_consist_ interaction are the only significant predictors. Segment interaction accounts for the largest relative effect (33.45%). Training *R*² = 0.19 suggests potential overfitting. Dashes (-) indicate statistics not applicable for that parameter type. | | | | | | | | | | | | | |

**Table S9.** Bivariate Segmented Regression Results (PFOS)

| **Component** | **Parameter** | **Value** | **SE** | ***t*** | ***p*** | **Training** | **Testing** | **% Effect** | ***n*** | **Mean** | **SD** | **Min** | **Max** |
| --- | --- | --- | --- | --- | --- | --- | --- | --- | --- | --- | --- | --- | --- |
| **Model Info** | Response Variable | Z_PFOS_ | - | - | - | - | - | - | - | - | - | - | - |
|  | Predictor 1 | δ^13^C_consist_ | - | - | - | - | - | - | - | - | - | - | - |
|  | Predictor 2 | δ^15^N_consist_ | - | - | - | - | - | - | - | - | - | - | - |
|  | Training Observations | 89 | - | - | - | - | - | - | - | - | - | - | - |
|  | Testing Observations | 23 | - | - | - | - | - | - | - | - | - | - | - |
| **Performance** | *R*² | - | - | - | - | 0.35 | -0.05 | - | - | - | - | - | - |
|  | MSE | - | - | - | - | 0.63 | 1.09 | - | - | - | - | - | - |
|  | RMSE | - | - | - | - | 0.79 | 1.04 | - | - | - | - | - | - |
| **Breakpoints** | δ^13^C_consist_ | 0.19 | - | - | - | - | - | - | - | - | - | - | - |
|  | δ^13^C_consist_ Below | 80 (71.4%) | - | - | - | - | - | - | - | - | - | - | - |
|  | δ^13^C_consist_ Above | 32 (28.6%) | - | - | - | - | - | - | - | - | - | - | - |
|  | δ^15^N_consist_ | 0 | - | - | - | - | - | - | - | - | - | - | - |
|  | δ^15^N_consist_ Below | 71 (63.4%) | - | - | - | - | - | - | - | - | - | - | - |
|  | δ^15^N_consist_ Above | 41 (36.6%) | - | - | - | - | - | - | - | - | - | - | - |
| **Coefficients** | Intercept | -0.61 | - | - | - | - | - | - | - | - | - | - | - |
|  | δ^13^C_consist_ | -0.45 | 0.2 | -2.8 | **0.01**** | - | - | 17.91 | - | - | - | - | - |
|  | δ^13^C_consist_ segment | 1.26 | 0.5 | 2.56 | **0.01*** | - | - | 50.55 | - | - | - | - | - |
|  | δ^15^N_consist_ | 0.03 | 0.2 | 0.19 | 0.85 | - | - | 1.1 | - | - | - | - | - |
|  | δ^15^N_consist_ segment | -0.05 | 0.3 | -0.2 | 0.86 | - | - | 1.93 | - | - | - | - | - |
|  | δ^13^C_consist_ × δ^15^N_consist_ interaction | 0.16 | 0.1 | 1.96 | 0.05 | - | - | 6.44 | - | - | - | - | - |
|  | Segment interaction | 0.55 | 0.7 | 0.85 | 0.4 | - | - | 22.06 | - | - | - | - | - |
| **Segment Combinations** | Low δ^13^C_consist_, Low δ^15^N_consist_ | - | - | - | - | - | - | - | 65 | -0.28 | 1 | -2.3 | 2.9 |
|  | Low δ δ^13^C_consist_, High δ^15^N_consist_ | - | - | - | - | - | - | - | 15 | -0.4 | 0.6 | -1.8 | 0.5 |
|  | High δ^13^C_consist_, Low δ^15^N_consist_ | - | - | - | - | - | - | - | 6 | 0.54 | 0.7 | -0.8 | 1.4 |
|  | High δ^13^C_consist_, High δ^15^N_consist_ | - | - | - | - | - | - | - | 26 | 0.8 | 0.9 | -0.4 | 3.4 |
| **Note:** Bold *p*-values indicate statistical significance (*p* < 0.05). Asterisks denote significance level (* *p* < 0.05, ** *p* < 0.01). The model shows better training performance than PFCA (*R*² = 0.35) but still poor testing performance (*R*² = -0.05). δ^13^C_consist_ and its segment term are highly significant. δ^13^C_consist_ segment accounts for the largest relative effect (50.55%). High δ^13^C_consist_ segments show positive means regardless of δ^15^N_consist_ levels. Dashes (-) indicate statistics not applicable for that parameter type. | | | | | | | | | | | | | |
